# Stink bug *Agonoscelis* spp. (Heteroptera: Pentatomidae)-An Emerging Threat for Seed Production in Alfalfa Crop (*Medicago sativa* L.) and Their Successful Management

**DOI:** 10.1101/2021.02.17.431601

**Authors:** Haider Karar, Muhammad Amjad Bashir, Abdul Khaliq, Muhammad Jaffar Ali, Reem Atalla Alajmi, Dina M. Metwally

## Abstract

Forages are vital constituent for sustainable agriculture because they provide feed for animals that ultimately converted into human food. Alfalfa is one of the most important forages that has highest feeding value for livestock, and seed production of alfalfa seriously affected by several factors, but seed yield loss due to stink bug attack is more as compared to other factors. Studies were conducted to control stink bug by different insecticides at Fodder Research Institute, Sargodha, Punjab-Pakistan during 2016-17. The efficacy of ten insecticides viz., acephate, dimethoate, malathion, chlorpyriphos, bifenthrin, lambdacyhalothrin, deltamethrin, acetamiprid, imidacloprid and carbosulfan were tested against stink bug, *Agonoscelis* spp. (Heteroptera Pentatomidae) on alfalfa, *Medicago sativa* L. variety SGD-2002. The mortality of stink bug was recorded one, three, five, seven, ten and fifteen days after spray. Similarly the population of pollinators was recorded before and one, three and five days after spray. From this study it had been observed that acetamiprid (81.14 %) and acephate (80.65%) had the higest mortality of stink bug and proved to be the most effective insecticides against stink bug. By spray of insecticides the population of pollinators declined one day after spray, but it had been rehabilitated three days after spray. By chemical treatment against stink bug, seed yield increased from 28.05 Kg/acre (during last four year without chemical control of stink bug) to 116 Kg/acre in 2016-17 (with chemical control). From this study it has been concluded that chemicals can be used in integrated management program of alfalfa seed production.

## 1. Introduction

Forages are considered pillar of sustainable agriculture because they provide feed for livestock which convert fiber into human food, such as milk and meat. Among forages, legumes play vital role worldwide in grassland ecosystem (Singh *et al*.2010). Alfalfa is one of the legumes forage which has highest feeding value for livestock due to its nutritious and quality, having 16-25% crude protein and 20-30% fiber (Babu *et al*., 2010). Alfalfa is considered as a rich source of minerals because 100 grams of alfalfa contains Potassium 79 mg, Phosphorus 70 mg, Calcium 32 mg, Magnesium 27 mg, Sodium 6 mg, Iron, Zinc and Manganese in traces. In Pakistan, the fodder and seed yield of alfalfa is low as compared to other developing countries. Many factors involved in yield reduction such as poor field management, unjustifiable last cutting, changing climatic conditions and poor management of insets, pests and diseases. A varieties of insects and pests attacked on alfalfa crop, resulted a dramtic decrease in yield and quality of produce (Godfrey *et. al*., 2013). Among sucking pests stink bug is considered as one of the most destructive pest of alfalfa at reproductive stage. Stink bug is considered one of the major pests of economically important crops worldwide (Panizzi, 1997), and it is also responsible for transmission of various diseases in plant (Medrano *et al*., 2009). The population of stink bug increased when plant reached to mature stage of pod formation. It sucks sap from the developing pods and seed formation seriously affected. It is highly polyphagous pest (Hoebeke and Carter, 2003) and may cause tremendous economic damage to crops worldwide. Stink bug is a mobile insect and can travel from nearby agricultural and wild hosts into farm; this movement linked to crop phenology and the availability of suitable food sources (Jones and Sullivan, 1982). It is not only notorious pest of alfalfa pods but it has more than 60 host plants (Bernon *et al*. 2004). According to an estimate, globally the annual crop losses are more due to stink bugs alone. For example, in cotton losses are $31 million (Williams, 2009) and in soybean $60 million (McPherson and McPherson, 2000). Similarly in grain and legume crops stink bug feeding declines the quality of the produce like seeds or bean value and whole heads or fruiting bodies can be lost (Hall and Teetes, 1982; Espino and Way, 2008). It is not only pest of legume crop but it is also considered as severe pests in Maize (Negron and Riley, 1987; Ni *et al*., 2010) and peach and can cause 100% crop losses (Leskey *et al*. 2012a). According to Wheeler 2001, suckling by alfalfa bug, *A. lineolatus* can decrease up to 50% or more seed yield while Sekulić *et al*. (2005) reported that the alfalfa plant bug reduced the seed yields of about 20-90 %.

Pollinators also play an important role in seed production. So there is a need to develop effective integrated pest management programs to reduce major crop losses. Stink bug is a recently noticed pest in Pakistan, so there is scarcity of cultural and biological strategies which can be helpful to develop sustainable pest management programs. This has resulted in the need for immediate insecticide based management program for the control of stink bug, *H. halys* (Leskey *et al*. 2012b) on various crops while other long term strategies can be developed for maximum alfalfa seed production. The insecticides are considered the only immediate effective option available for producers to minimize economic losses.

Alfalfa is a 100 percent cross pollinated crop and its optimum seed production depends upon the activity of pollinators. Among pollinators bees play an important role in fertilization of flowers. It is necessary to screen different groups of insecticides with different modes of action, to evaluate the chemicals which ensure minimum damage to pollinators for maximum seed production of alfalfa seed crop. The fodder crops are an integral part of livestock management. Fodder crops although attacked by various pests but are seldom treated with pesticides in course of its management because the crops are direct feed of livestock. But, in case of seed production, the fodder crops are kept in the field for flowering and seed formation and in this case the crop is not fed to livestock. The chemical treatment of fodder crops for seed production purpose not only enhances seed production but it is relatively safe because it is not used as feed for livestock.

The purpose of this study was to control the stink bug through that insecticide which are safe for natural pollinators such as honey bees for the enhancement of seed production.

## 2. Materials and Methods

The experiments were conducted at Fodder Research Institute, Sargodha-Punjab Pakistan during 2016-17. Detail of different insecticides used in this study is given in Table 1. One Alfalfa variety was used in this study, last cutting left to obtain seed. Doses of different insecticide were used in water solution and applied by manually operated hand knapsack sprayer at the rate of 100 liter solution acre ^-1^ in plots of size 10×15m. The experiment was arranged in randomized complete block design (RCBD) with three replications. The data of stink bug was recorded before spray, then 1, 3, 5, 7, 10 and 15 days after spray by the use of ten net sweeps per plot. All insecticides were sprayed at 6.00 PM to save pollinators population. The data regarding activity of pollinators were recorded at 10.00 AM before and then after one, three and five days spray. Before spray of each insecticide, spray machine was cleaned thoroughly with clean water to avoid insecticide mixture. The data was compiled and statistically analyzed.

### 2.1. Mortality rate

Percent mortality of stink bug was calculated by using following formula:

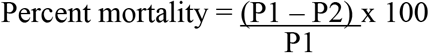

Population before spray= P1

Population after spray=P2

### 2.2. Pollinators

The population of pollinator was recorded from alfalfa seed crop from five different places. The handmade square meter was kept in the field and counted the population of all visiting insects. Average pollinators populations were calculated per square meter.

### 2.3. Yield data

The crop was harvested in the month of June 2017 and kept on concrete floor for drying. After three days the crops was thrashed and yield was recorded. The yield data of last four years was collected from farm manager, Fodder Research Institute, Sargodha and comparative graph was made.

### 2.4. Statistical analysis

All the treatments were compared with control and one another to assess the performance of the treatment in uncontrolled field condition. The data were finally subjected to statistical analysis using Statistix version-9 (www.statistix.com/freetrial.html) and means were compared by Tukey’s HSD.

## 3. Results

### 3.1. Mortality rate one day after spray

The effect of various insecticides on mortality rate of stink bug population was presented in Table 2. There was a significant difference (P<0.01) among treatments. Carbosulfan and bifenthrin sprayed plots showed 95.65% and 84.62% mortality of the pest that are statistically similar with deltamethrin, imidacloprid, dimethoate, chlorpyrifos, and acetamiprid while malathion sprayed plots showed 52.63% morality of the pest.

### 3.2. Mortality rate three days after spray

The effect of different pesticides on mortality rate of stink bug population three days after spray was shown in Table 2. Three days after pesticides spray acephate and acetamiprid sprayed plots show 100% mortality rate of stink bug which are statically similar with imidacloprid and carbosulfan that showed mortality of stink bug 80.0% and 75.46% respectively. Lambdacyhalothrin and bifenthrin sprayed plots showed 29.55% and 23.21% respectively, mortality rate of stinkbug three days after spray.

### 3.3. Mortality rate five days after spray

Significant differences (P<0.01) in stink bug population were observed among different treatments five days after spray. Five days after spray acephate maintained its highest mortality rate of stink bug population (100%), followed by acetamiprid (82.76%) and carbosulfan (52.61%), while in plots where lambdacyhalothrin and bifenthrin were sprayed the mortality rate of stinkbug was zero.

### 3.4. Mortality rate seven days after spray

Significant variations (P<0.01) was observed among different treatments. Acephate (84.62%) and acetamiprid (81.82%) sprayed plot maintain highest mortality rate of stink bug as compared to other insecticides seven days after spray, while lambdacyhalothrin, deltamethrin, dimethoate, malathion and bifenthrin sprayed plots show lowest mortality rate of stink bug (Table 2).

### 3.5. Mortality rate ten days after spray

The effect of different insecticides on mortality rate of stink bug population ten days after spray was shown in Table 2. Highest mortality of stink bug was recorded in acetamiprid (81.82%) and acephate (68.56%) sprayed plots, while in plots where lambdacyhalothrin, chlorpyrifos, deltamethrin, dimethoate, malathion and bifenthrin sprayed the mortality of stink bug was zero ten days after spray.

### 3.6. Mortality rate fifteen days after spray

The data regarding percent mortality of the pest revealed significant variations (P<0.01) among different treatments. The maximum mortality of stink bug was observed in acetamiprid (67.73%) and acephate (61.54%) sprayed plots fifteen days after spray. Whereas, 0.00 percent mortality of the pest was observed in those treatment where lambdacyhalothrin, chlorpyrifos, deltamethrin, dimethoate, malathion and bifenthrin were sprayed fifteen days after spray.

### 3.7. Cumulative mortality of stink bug

The data regarding stink bug mortality in alfalfa seed crop on cumulative basis are graphically shown Fig. 1. The results revealed that mortality of stink bug in acephate and acetamiprid spray plots was 80.65% and 81.14% respectively, while the mortality of stink bug in other insecticides sprays plots ranged from 16.59% to 44.35%.

### 3.8. Position of pollinators before and after spray

The effect of different insecticides spray on pollinators population in alfa crop is shown in Fig. 2. All insecticides spray significantly reduced the population pollinators one day after spray, the highest and lowest reduction of pollinators population was observed in chlorpyrifos and acetamiprids sprayed plots respectively, one day after spray. However, negligible effect of insecticides spray on pollinators population was observed three days after spray, because the population of pollinators rehabilitated three days after spray (Fig. 2).

### 3.9. Seed yield of Alfalfa during last five years

Seed yield of Alfalfa crop and population of stinkbug during last five years are shown in Fig 2. Inverse relationship between seed yield and stinkbug population had been observed during various years, seed yield of Alfalfa crop in 2012-13, 2013-14, 2014-15, 2015-16 and 2016-17 was 42.0, 10.5, 22.5, 37.2 and 116.0 Kg acre ^-1^ respectively, while stinkbug population was 4.19, 5.93, 6.09, 4.74 and 0.02 individuals per net sweep respectively.

## 4. Discussion

Production of alfalfa seed depends upon several factors i.e. last cutting dates, weather factors, availability of pollinators, insect pests and diseases. Among these yield limiting factors, insect pests are considered the utmost important factor which need special attention. Different management practices are being adopted globally viz., cultural, biological and chemical to overcome insect pests of various crops. But the success of pest control is judged by results and the best control strategy that gives adequate pest control. Among these control measures chemical control is considered to be the most effective and efficient strategy (Kuhar *et. al*. 2015b) which saves the crop from pest outbreak. In our experiment, four different groups of insecticides with different mode of action were tested against stink bug on alfalfa seed crop under field conditions. Our results suggested that all insecticides were statistically different and had significant impact on stink bug mortality when compared with control plot. On numerical basis, maximum percent mortality was observed in acetamiprid, acephate and proved most effective insecticide as compared with other insecticides (Table 2). The results are in agreed with that of Karar and Khaliq, 2019 who concluded that acetamiprid, acephate proved to be the best insecticides against seed pods feeding stink bug with increase in yield. Further it was noted that carbosulfan showed knock down effects and remained effective for short period against stink bug. Similar results reported by Wallingford (2012) and Lee *et al*. (2013a) who said that organophosphates, neonicotinoids, pyrethroids and carbamates insecticides have been the utmost effective and efficient strategy to manage stink bug. Similarly Leskey *et. al*. (2012) described that acetamiprid showed greatest mortality of stink bug i.e. 93-100%. Furthermore our results supported by Leseky (2014) who partially confirmed that pyrethroids, organophosphate, acephate, carbamates, methomyl and oxamyl, neonicotinoids dinotefuran, imidacloprid, thiamethaxim, clothianidin, and acetamiprid were also effective to control stink bug. In our study it has been observed that pyrethroids like bifenthrin, lambdacyhalothrin and deltamethrin gave highest mortality of stink bug one day after spray but after three days the mortality reduced, that are in line with the study of Leskey *et. al*. (2012) who reported that pyrethroids have knockdown effects but many bugs recover within 7 days. Similarly in our results carbamate like carbosulfan has also knock down effect and remained effective for period of five days.

Seed yield of crop and stink bug population had inverse relationship, when stink bug population increased the seed yield of Alfalfa dramatically decreased (Fig 3). When stink bug population controlled by insecticides there was a multiple times increased in seed yield. This is first study in Pakistan that indentified the stink bug population dramatically reduced seed yield of Alfalfa crop and its control by insecticides dramatically increased the seed yield of Alfalfa crop. The alfalfa crop is highly cross pollinated. For the production of seed it is important to enhance the activity of pollinators for more seed production. It had been observed that insecticides spray had negligible effect on the population of pollinators because population of pollinators rehabilitated three days after spray

Comparison of alfa alfa seed production in 2016-17 with last four years data clearly depicts that maximum yield of alfalfa seed was obtained during 2016-17 (FRI-Sargodha) as compared to last four years yeild. The data shows that there is much gap between previous years and current year yield. The reasons could be that there is lack of knowledge regarding detrimental effect of stink bug on seed yield of crop and importance of pollinators activity to enhanced seed yield. When the pest was properly managed and increased the activity of bees the yields boost up multiple times. Anyhow, further studies should also be carried out to find suitable selective insecticides which ensure maximum mortality of stink bug and minimum mortality of pollinators.

## 5. Conclusions and Recommendations

Stink Bug *Agonoscelis* spp. (Heteroptera: Pentatomidae) which was seriously noticed as a pest in alfalfa seed crop during 2016-17 in Sargodha-Punjab, Pakistan. Increase in the activity of pollinators and management of stink bug play a key role in increasing the alfalfa seeds production. It is expected that it may be warning for other economically important agricultural crops like citrus, grapes, apple, fig, peach, pear, cotton, tomato, corn and soybeans in future. This study generates awareness among growers to focus their attention on cultivated crops regarding newly emerged insect pest. The present project indicates that acetamiprid and acephate were the most effective insecticides for the management of stink bug as compared to other insecticides tested. Carbosulfan was also effective to some extent for short period of time, with knock down effect. These insecticides have diverse mode of action and can be included in the IPM module which will be very helpful in planning future program of pest management and keep them as part of chemical control to avoid resistance and cross-resistance, which ultimately result in better and healthy crop production and increase the GDP of the country. It is further recommended that spray should be done after 6.00 pm to save and increased activity of pollinators in seed production.

## Acknowledgement

Current studies were conducted at Fodder Research Institute, Sargodha - Punjab, Pakistan with the coordination of Fodder Botanists, Malik Amir Abdullah and Mr. Ghulam Nabi, and Abdul Razzaq, Assistant Soil Chemist, Government of the Punjab, Pakistan. Authors are thankful to our laboratory Assistant, especially M. Nasrullah and other field staff of Fodder Research Institute, Sargodha for assistance in present studies.

## Author’s Contributions

HK developed idea and layout of the study. HK wrote up the manuscript. NM reviewed and rephrased the manuscript. MJA statistically analysis, AK collected data and MH,NH help in preparation of manuscript.

